# Large conformational changes in FtsH create an opening for substrate entry

**DOI:** 10.1101/209445

**Authors:** Vanessa Carvalho, Roland Kieffer, Nick de Lange, Andreas Engel, Marie-Eve Aubin-Tam

## Abstract

AAA+ proteases are degradation machines, which exploit ATP hydrolysis to unfold protein substrates and translocate them through a central pore towards a degradation chamber. FtsH, a bacterial membrane-anchored AAA+ protease, plays a vital role in membrane protein quality control. Although cytoplasmic structures are described, the full-length structure of bacterial FtsH is unknown, and the route by which substrates reach the central pore remains unclear. We use electron microscopy to determine the 3D map of the full-length *Aquifex aeolicus* FtsH hexamer. Moreover, detergent solubilisation induces the formation of fully active FtsH dodecamers, which consist of two FtsH hexamers in a single detergent micelle. FtsH structures reveal that the cytosolic domain can tilt with respect to the membrane. A flexible linker of ~20 residues between the second transmembrane helix and the cytosolic domain permits the observed large tilting movements, thereby facilitating the entry of substrate proteins towards the central pore for translocation.

## Introduction

Cells are complex systems that rely on numerous tightly controlled vital processes. For instance, protein quality control is crucial to the maintenance of the cell’s proteome. To avoid the lethal accumulation of misfolded or non-functional proteins, eukaryotes as well as prokaryotes use proteolysis (Barrett et al., 2012). In this process, peptide bonds are cleaved by proteases, and resulting amino acids (aa) are reused to build new and functional proteins. This cycle allows cells to maintain their homeostasis and to keep them healthy. It is then understandable that malfunctions in proteolysis lead to diverse forms of diseases (López-Otím and Bond, 2008; Richard, 2005).

AAA+ proteases belong to the family of ATPases associated with various cellular activities, and are molecular machines capable of unfolding and degrading proteins (Olivares et al., 2016). AAA+ proteases share several structural and functional characteristics. They assemble into a barrel-shaped chamber with a central pore formed by the ATP-binding domains. The pore entrance exhibits translocating loops with highly conserved residues, which bind to target substrates. ATP-driven conformational changes of the ATP-binding domain unfold the bound substrate and translocate it through the central pore into the proteolytic chamber for degradation. In general, bacterial AAA+ protease malfunctions can lead to a complete discoordination of the cell homeostasis.

From the five AAA+ proteases in *E. Coli*, FtsH is the only one that is anchored to the membrane and that is essential (Bittner et al., 2017). FtsH plays a crucial role in membrane protein quality control (Hari and Sauer, 2016), and in aminoglycoside antibiotic resistance, possibly by eliminating misfolded proteins disruptive to the membrane (Hinz et al., 2011). FtsH also regulates the phospholipid to lipopolysaccharide (LPS) ratio in the outer membrane by degrading LpxC, the key enzyme of LPS biosynthesis (Schäkermann et al., 2013).

In mitochondria, the i-AAA protease, a FtsH ortholog, translocates polynucleotide phosphorylase into the intermembrane space (Rainey et al., 2006), while the hydrophobicity of a specific transmembrane segment dictates its dislocation from the inner membrane by the mitochondrial *m*-AAA protease, another FtsH ortholog (Botelho et al., 2013). In humans, mutations in the gene coding for paraplegin, a subunit of *m*-AAA, are related to the severe disease *spastic paraplegia* (Nolden et al., 2005). Therefore, and because the FtsH mechanics and structure are less well understood than those of cytoplasmic AAA+ proteases, increasing our knowledge on the workings of FtsH are of both medical and fundamental interest.

The FtsH protein comprises a N-terminal transmembrane helix, a ~75 aa periplasmic domain, a second transmembrane helix (Scharfenberg et al., 2015), and the larger cytoplasmic AAA+ and protease domains (Bieniossek et al., 2009). FtsH proteins assemble into hexamers, with 12 transmembrane helices inserting into the lipid bilayer. The ATPase domain has conserved arginine residues that compose the second region of homology (SRH), which is believed to be crucial for FtsH oligomerization. This domain also houses the highly conserved Walker A and Walker B domains, which bind and hydrolyse nucleotides (Bieniossek et al., 2009; Vostrukhina et al., 2015).

Structural studies have used truncated FtsH forms with only the soluble C-terminal (cytosolic) part (Bieniossek et al., 2009; Bieniossek et al., 2006; Kim et al., 2008; Niwa et al., 2002; Suno et al., 2006; Suno et al., 2012; Vostrukhina et al., 2015) or with only the periplasmic domain (Scharfenberg et al., 2015). The single full-length structure concerns *m*-AAA, the yeast mitochondrial ortholog of bacterial FtsH, which has been resolved at 12 Å resolution by cryo-electron microscopy (Lee et al., 2011). Therefore, no information on the conformational rearrangement of full-length FtsH in relation to the membrane when bound to nucleotides or to a substrate is available. Crystal structures of the cytosolic domain of FtsH exhibit a six-(Bieniossek et al., 2009), a two-(Bieniossek et al., 2006; Vostrukhina et al., 2015) or a three-fold (Suno et al., 2006) symmetric conformation of the ATPase domain. These different conformations suggest that the ATPase domain could move polypeptides in steps as long as 45 Å into the central cavity during ATP hydrolysis cycles (Bieniossek et al., 2009). In contrast, the C-terminal protease domain always shows a six-fold symmetry for all crystal structures, i.e., the cytosolic domain of *Thermus thermophiles* FtsH (Suno et al., 2012), of *Thermotoga maritima* FtsH (Bieniossek et al., 2009; Bieniossek et al., 2006) and of *Aquifex aeolicus* FtsH (Suno et al., 2006; Vostrukhina et al., 2015). The proposed mechanism for substrate entry in *m*-AAA is based on substrate recognition by solvent exposed lateral regions of FtsH cytosolic domain. Accordingly, a 13 Å gap between the membrane and the cytosolic domain observed by cryo-electron microscopy would provide access to substrate, which implies that only (partly) unfolded proteins can reach the translocating loops and be moved through the pore for degradation (Lee et al., 2011).

Here we report the first full-length structure of *Aquifex aeolicus* FtsH (AaFtsH), which we determined by negative stain electron microscopy to a resolution of 20 Å. Unexpectedly, upon detergent solubilisation not only hexamers of AaFtsH formed, but also fully stable and active dodecamers of AaFtsH. The dodecamer structure is solved to a resolution of 25 Å, showing two AaFtsH hexamers sharing a single Lauryl Maltoside Neopentyl Glycol (LMNG) micelle, with the periplasmic domain of one hexamer touching the cytosolic domain of the other hexamer. This arrangement induces a tilt of the periplasmic domain with respect to the cytoplasmic domain. While AaFtsH hexamers and dodecamers have similar ATPase and proteolytic activity, the latter exhibit a substantial larger gap between the cytosolic domain and the transmembrane domain. Therefore, we propose that the cytosolic domain can tilt with respect to the membrane plane for substrates to reach the translocating pore loops, as required for substrate unfolding and degradation.

## Results

### Full-length AaFtsH purification

AaFtsH with a C-terminal His-tag was overexpressed in *E.Coli* cells and extracted from purified membranes with the use of a mild detergent, LMNG (Chae et al., 2010). Ni-NTA chromatography followed by size-exclusion chromatography (SEC) using Superose 6 10/300 GL yielded pure AaFtsH complexes (Figure 1). Best results were obtained by incubating the AaFtsH containing fractions, collected from Ni-NTA chromatography, overnight at 60°C in presence of 20 mM ATP, 10 mM MgCl_2_ and 25 μM ZnCl_2_ just before the SEC purification. The SEC profile shows a first peak that is centred at 12.1 mL ± 0.2 mL (SD, N=10) and a second peak at 13.4 mL ± 0.2 mL (SD, N=10) (Figure 1A). Peak positions are determined by simultaneously fitting two Gaussian functions. Native gel electrophoresis suggests that the second peak likely contains hexameric AaFtsH as it runs at ~700 kDa, while the first peak indicates a higher oligomeric state (Figure 1B). The molecular weight of the eluted complexes was estimated using the partition coefficient (K_av_) values extracted from a calibration curve of the Superose 6 column, and the positions of fitted Gauss functions. The second peak is centred at a molecular weight of 730 kDa, which is larger than the size expected for AaFtsH hexamers (~432 kDa). Extrapolating the calibration curve to smaller elution volumes, the first peak corresponds to a molecular weight of 940 kDa.

**Figure 1.**
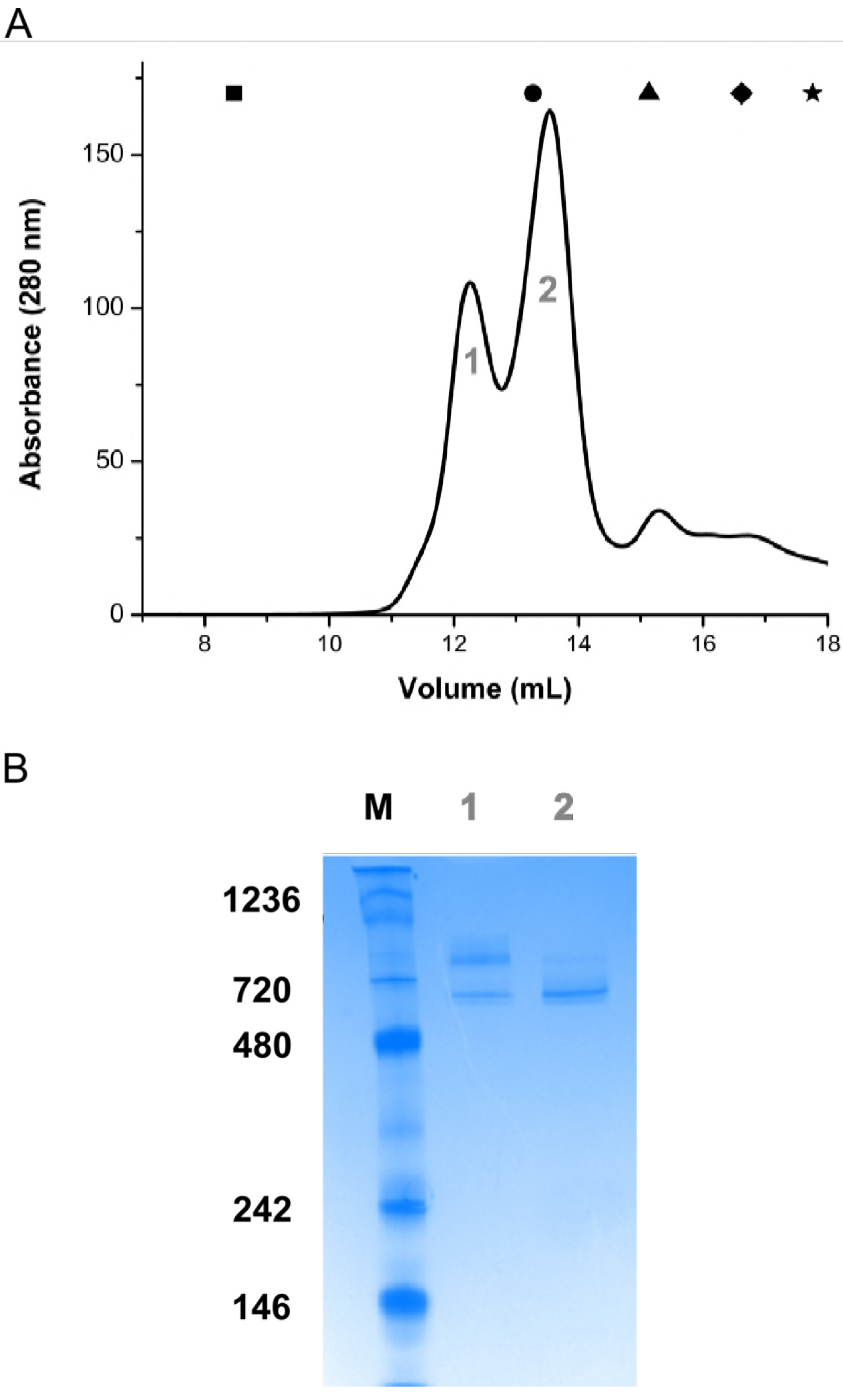
FtsH oligomers elute in two peaks from sizing chromatography. *A*. Representative size exclusion chromatography run. Blue Dextran (◾) used to calculate the void volume of the column (8.47mL). The elution volumes of standard proteins used to estimate the molecular weight of the eluted fractions are shown on the top of the graph: (•) Tyroglobulin (MW: 669 kDa; V_e_: 13.27 ml), (▲) Ferritin (MW: 440 kDa; V_e_: 15.13 ml), (♦) Aldolase (MW: 158 kDa; V_e_: 16.62 ml) and (★) Ovalbumin (MW: 44kDa; V_e_: 17.76 ml). B. Native PAGE gel of peak 1 and 2 (lanes 1 and 2) and molecular marker (M).

### Electron microscopy of full-length AaFtsH complexes

Transmission electron microscopy (EM) visualized the difference of negatively stained samples from each peak. Preparations from the first peak revealed elongated particles with an average length of 231 Å ± 15 Å (SD, N=280; Figure 2A). Most particles from the second peak are smaller and exhibit an average length of 154 Å ± 13 Å (SD, N=280) (Figure 2B). Particles from the first peak appear to house two AaFtsH oligomers from the second peak, although their average length is less than twice the average length of the smaller particles.

**Figure 2.**
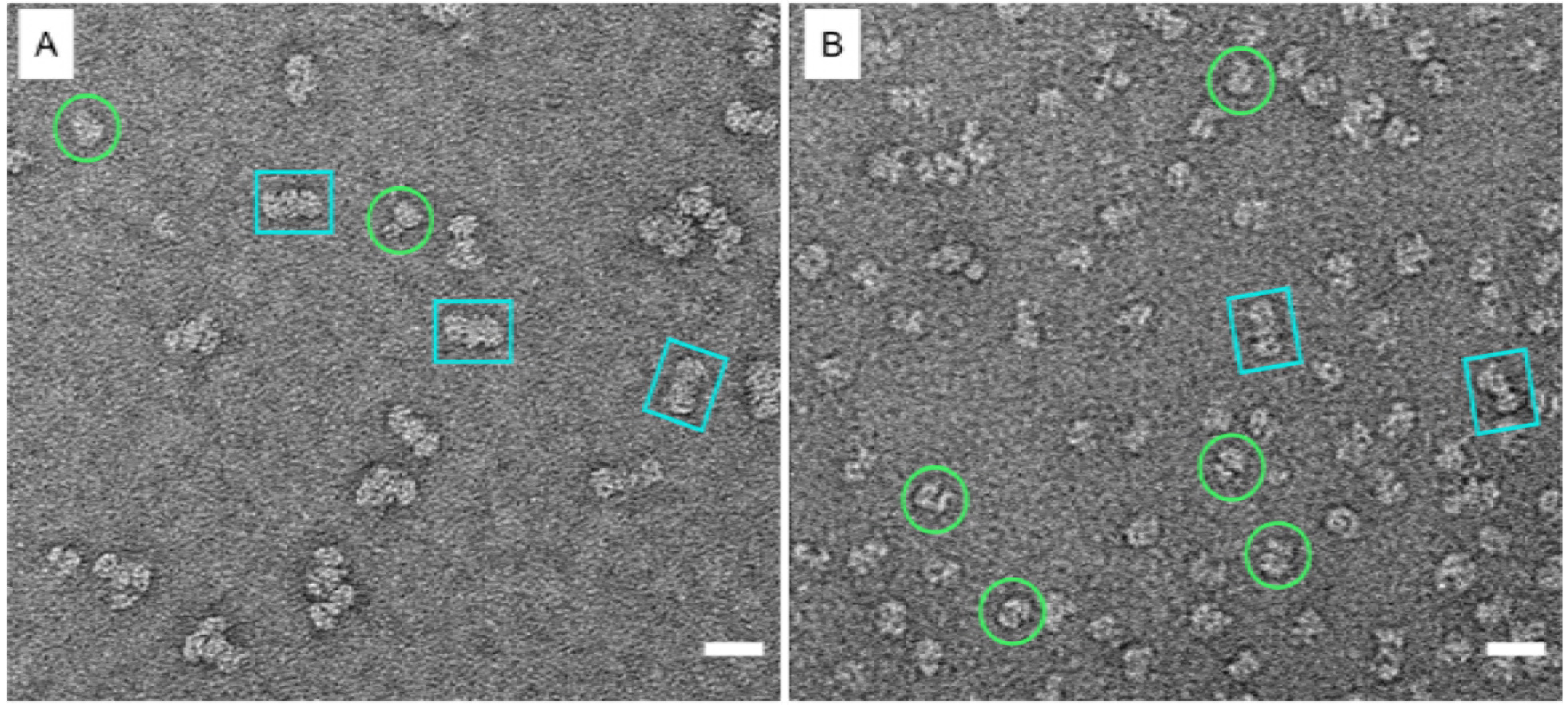
Micrographs of the first (*A*) and second (*B*) SEC eluted fractions prepared by negative staining. Side views are preferentially obtained. Blue squares correspond to dodecamers particles and green circles to hexamers. Scale bar is 200 Å.

### 2D Class averages and 3D reconstruction of the small AaFtsH complex

15000 projections of the smaller AaFtsH complex prepared by negative staining were selected and classified using the image processing software packages Scipion1.1 (Rosa-Trevín de la et al., 2016) and Eman2.12 (Ludtke, 2016). The class average corresponding to the infrequent, approximately axial projections is compatible with the hexameric nature of the small AaFtsH complex (Figure 3A), while Figure 3B-D display representative side or tilted views of AaFtsH hexamers. From such class averages EMAN2.12 calculated starting models and refined one of them against all particle projections, imposing 6-fold symmetry. Figure 3F displays the 20Å resolution 3D map of the AaFtsH hexamer, which accommodates the crystal structure of the AaFtsH cytosolic domain (PDB 4WW0) and the *E. coli* periplasmic domain (PDB 4V0B; Figure 3G).

**Figure 3.**
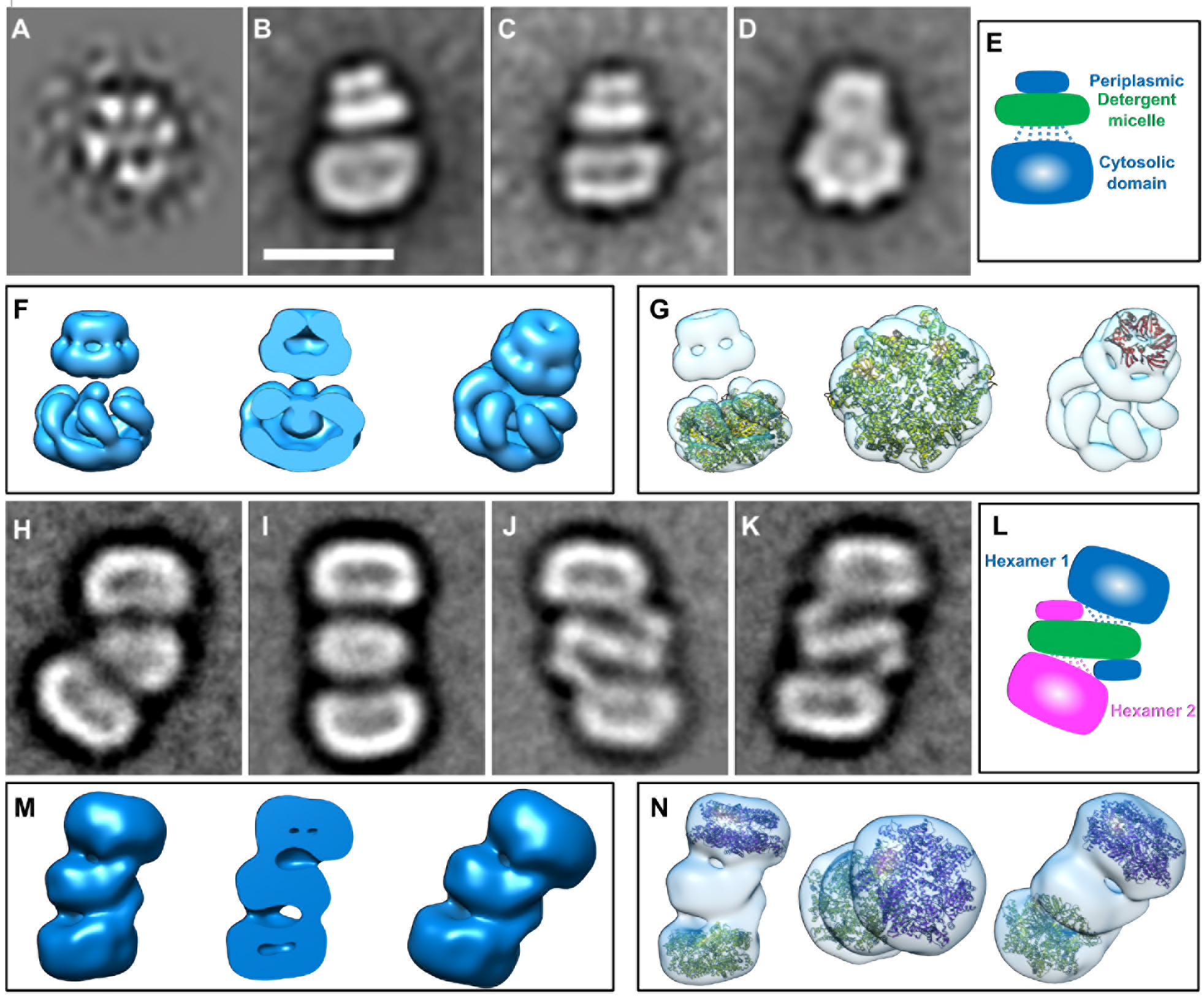
Class averages of negatively stained FtsH complexes and 3D maps. Class averages (*A-D*, *H-K*) represent top views (*A*), side views (*B* and *C*) and a view of a tilted hexamer (D), and different conformations of dodecamers (*H-K*).*E* and *L* – Schematic representation of the class averages highlighting with dashed lines six linking peptides present in the protein sequence. *F*, *M* – 3D maps of AaFtsH full length hexamer and dodecamer, respectively. *G* – Fitting of the crystal structure of the cytoplasmic domain of AaFtsH (PDB 4WW0) and the crystal structure of the periplasmic domain of *E. coli* FtsH (PDB 4V0B) to the hexamer map. *N* – Fitting of two cytoplasmic domains (PDB 4WW0) to the dodecamer map. Scale bar equals 110 Å.

### 2D Class averages of the large AaFtsH complex highlights large conformational changes

15000 projections of the larger elongated complex yielded class averages that document the conformational flexibility of AaFtsH dodecamers (Figure 3H, I). To obtain a 3D map of the dodecamers, class averages that correspond to a cylindrical (Figure 3I-K) rather than a bent conformation (Figure 3H) were selected to calculate an initial model that was subsequently refined against all projections of these classes without imposing any symmetry. Figure 3M and N illustrate the resulting 3D map and indicate how the structure of the AaFtsH cytosolic domain (PDB 4WW0) fit. As displayed in Figure 3L, class averages and 3D maps suggest that the AaFtsH dodecamer is kept in solution by a single LMNG micelle that embraces 2×12 transmembrane helices (green), while the periplasmic domains of each hexamer contact the cytosolic domains of the other hexamer (pink and blue).

### Dimensions of FtsH subunits and their conformational changes

Using ten class averages from both AaFtsH hexamers and dodecamers we measured that the detergent micelle, highlighted in green (Figure 3E,L), has a thickness of 40 ± 4 Å for the hexamer and 43 ± 2 Å for the dodecamer, which is close to the lipid bilayer thickness. The measured micelle/helical bundle width is 100 ± 18 Å for the hexamer and 126 ± 7 Å for the dodecamer. The estimated length of the 3D map for the full hexamer is 167 ± 5 Å (SD, N = 10) and its width is 131 ± 7 Å (SD, N = 10). When compared with the dimensions of *A. aeolicus* FtsH crystal structure, a similar width is reported (120 Å) (Suno et al., 2006). In hexamers, the cytoplasmic domain has a height of 83 ± 7 Å, and the periplasmic domain has a height of 31 ± 3 Å and a width of 63 ± 6 Å. The estimated length of the dodecamer 3D map is 243 ± 8 Å, nevertheless it accommodates two tilted and distorted hexamers. The height of the volume encompassing the micelle plus the two periplasmic domains is 90 ± 6 Å, which is comparable to twice the periplasmic domain (31 Å) plus the micelle height (40 Å). Albeit the medium resolutions achieved (25 Å for the dodecamer and 20 Å for the hexamers of AaFtsH), the orientation of the cytosolic domain with respect to the detergent micelle is clearly apparent. All the dodecamer classes and some of the hexamer classes show that the plane of the protease domain is at an angle relative to the detergent micelle creating a gap at the substrate entry site. This is more evident in the dodecamer conformations (Figure 3J-L), where a gap as large as 20 Å is observed between the central pore of AaFtsH cytosolic domain and the detergent micelle, and a gap of ~30 Å between the edge of that domain and the micelle (Figure 3 - figure supplement 1). This gap is large enough to accommodate the whole periplasmic domain of the second hexamer to which it is complexed.

**Figure 3 – figure supplement 1.**
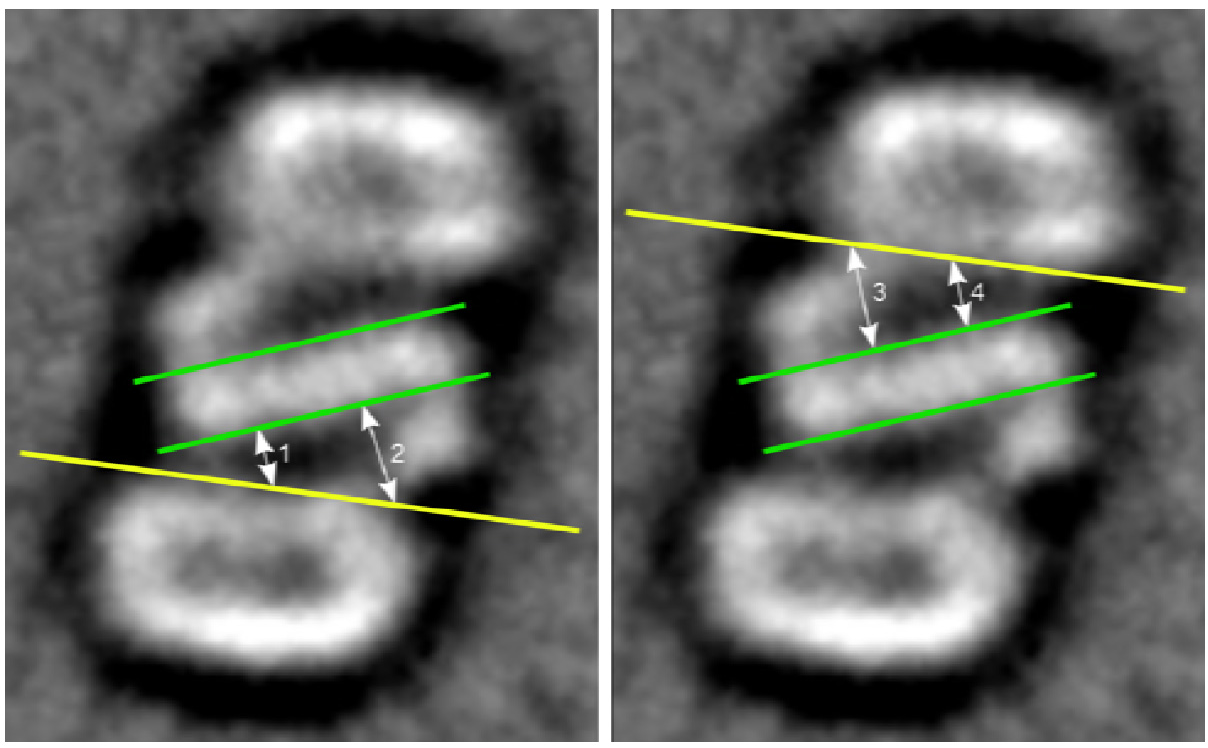
Measurements between the cytosolic domain and the detergent micelle. The distance between the central pore of AaFtsH cytosolic domain (delimited by the yellow line) and the detergent micelle (green lines) is approximately 20 Å (distances 1 and 4 are 18 Å and 20 Å, respectively), while ~30 Å (distances 2 and 3 are 32 Å and 33 Å, respectively) is measured between the edge of that domain and the micelle. This gap is large enough to accommodate partially folded proteins comparable in size to the periplasmic domain.

### Full-length AaFtsH hexamers and dodecamers show similar ATPase and protease activity

ATP hydrolysis rates were assessed by measuring the inorganic phosphate (Pi) released (Material & Methods). The initial velocity of the ATP hydrolysis reactions is extracted from the concentration of Pi released in the first 10-minutes interval (Figure 4, figure supplement 1). From this, specific ATPase activities of 338 nmol/min/mg and 340 nmol/min/mg were calculated for AaFtsH hexamers and dodecamers, respectively. To further compare the activity of FtsH in the hexameric and dodecameric forms, Michaelis Menten constants (K_M_ and K_cat_) were fitted using initial velocities calculated at various ATP concentrations (Figure 4A, B), with the use of FtsH monomer concentrations in the calculations (Bruckner et al., 2003; Tomoyasu et al., 1995). The results of the fits show that the dodecamer fraction has a K_M_ = 518 μM (95% Confidence Interval (CI): 432 - 625 μM) and a K_cat_ = 61 min^−1^ (95% CI: 57 − 66 min^−1^). The hexamer fraction has a K_M_ = 600 μM (95% CI: 480 − 754 μM) and a K_cat_ = 63 min^−1^ (95% CI: 58 − 69 min^−1^), identical to the values for the dodecameric AaFtsH within error limits. ATPase activity was measured at 60°C. The optimal growth temperature of *Aquifex aeolicus* is 85°C (Adams and Kelly, 1998); however, LMNG-solubilized AaFtsH aggregates at 80°C. To ensure that it is valid to perform the Malachite Green assay at 60°C, a control experiment was performed to calculate the measured release of Pi in absence of AaFtsH (Figure 4, figure supplement 1).

Next, we assessed the proteolytic activity of the hexamer and the dodecamer forms of AaFtsH. As previously reported (Akiyama, 2002; Vostrukhina et al., 2015), we used resorufin-labelled casein substrate (Roche) to test the proteolytic activity over a range of AaFtsH concentrations (Material & Methods). The initial rates of proteolytic activity were calculated from the concentration of resorufin released in the first 30-minutes interval (Figure 4, figure supplement 2). We assessed initial velocities for different AaFtsH monomer concentrations using the dodecamer (Figure 4C) and the hexamer (Figure 4D) fractions. The hexamers and the dodecamers reaction rates both increased linearly with AaFtsH concentrations at a slope of ~1.8 nM of resorufin per minute per μM of AaFtsH. This validates the idea that both fractions have the same proteolytical activity, suggesting that their active centres are not only well folded, but also equally accessible for protein entrance.

**Figure 4.**
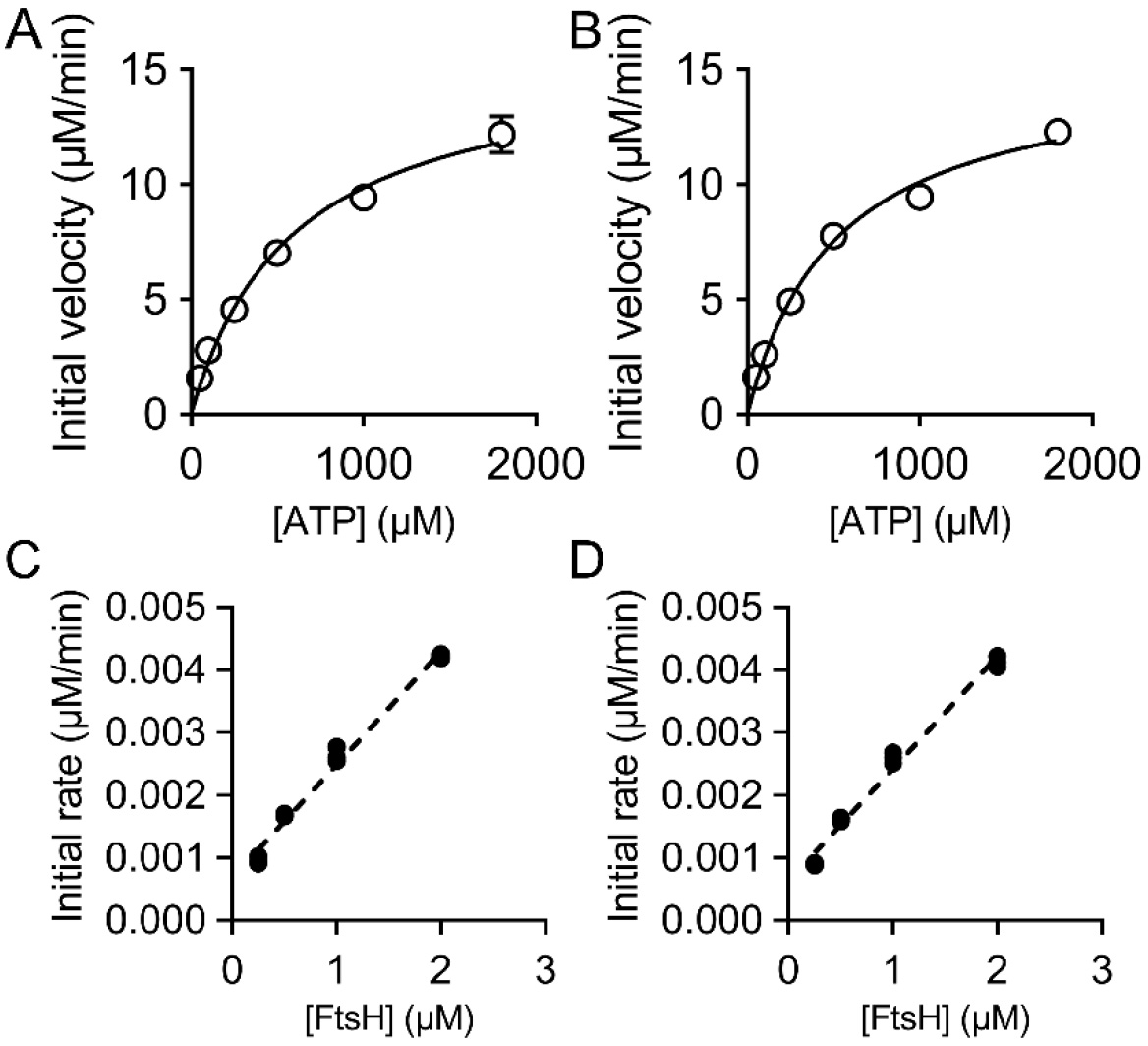
ATPase and protease activity assays. *A, B* – Steady-state velocities for the ATP hydrolysis by 0.25 μM AaFtsH dodecamers (*A*) and hexamers (*B*). The average and SD of three replicas for each reaction are plotted (SD bars are not visible for some points due to their small sizes). *C, D* – Initial velocities of proteolysis of 50 μM resorufin-labelled casein by AaFtsH dodecamers (*C*) and hexamers (*D*) as a function of AaFtsH concentration. All the reactions were measured three times and all points are plotted.

**Figure 4 – figure supplement 1.**
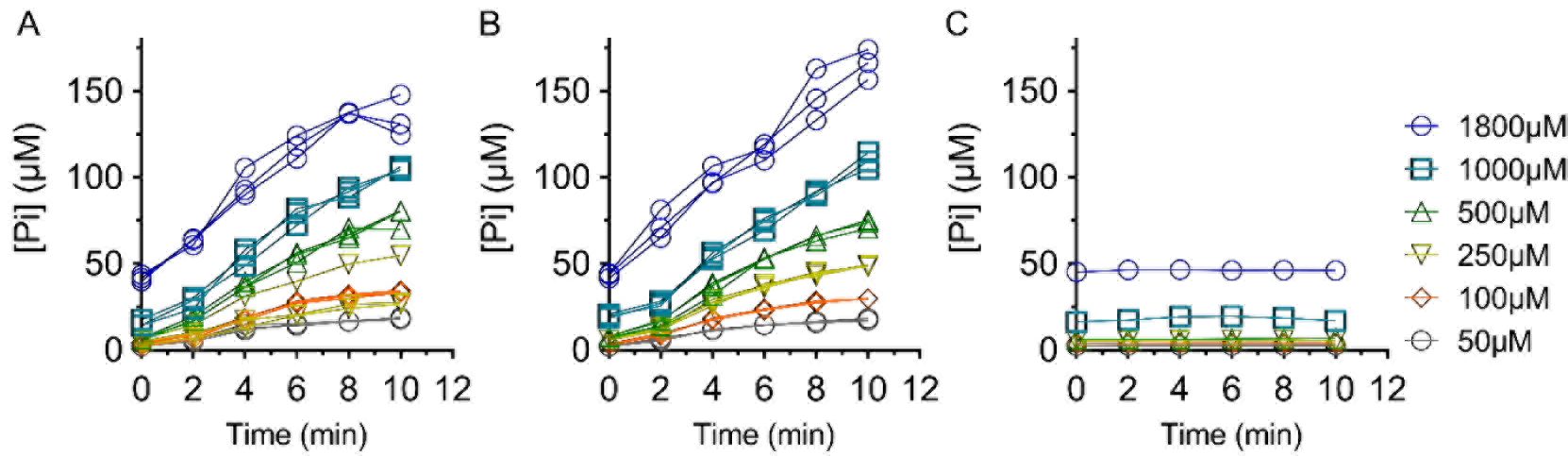
Raw data for the inorganic phosphate release. Free phosphate was measured using the Malachite Green assay kit for 0.25μM of AaFtsH dodecamer (*A*) and hexamer (*B*) fractions incubated with different ATP concentrations for 10 min. *C* – The free phosphate measurements in absence of AaFtsH.

**Figure 4 – figure supplement 2.**
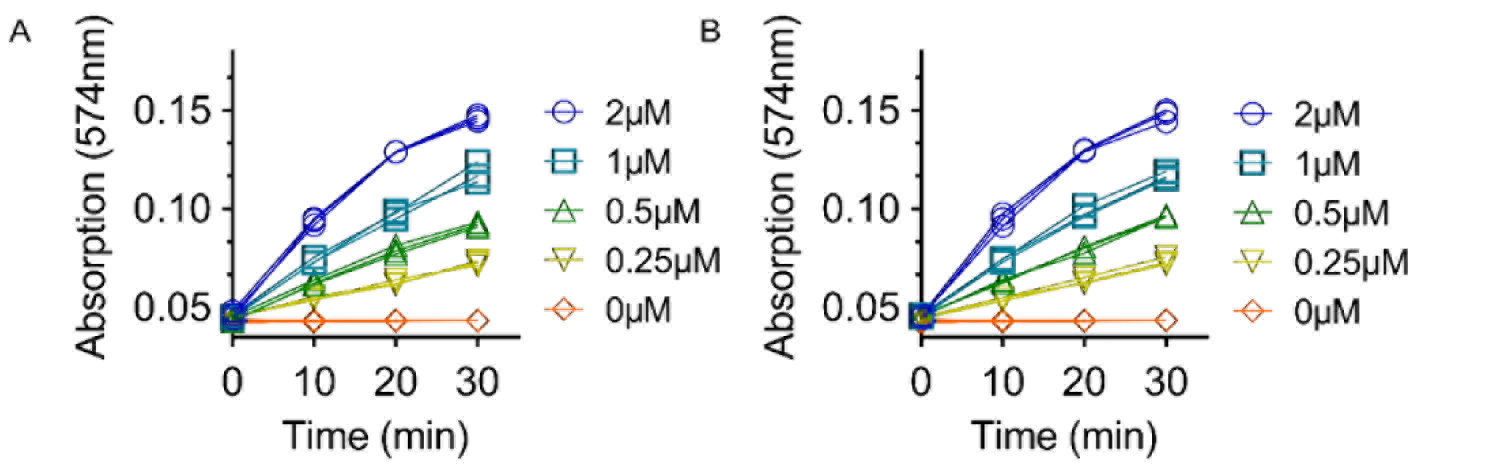
Protease activity assays raw data. Resorufin release was measured over 30 minutes for 50μM of Resorufin labelled casein incubated with dodecamer (*A*) and hexamer fractions (*B*) at different concentrations.

### Bioinformatics tools identify a linker region of ~20aa

To gain a better understanding of the unexpected conformational flexibility of AaFtsH complexes, we compared the *E. coli* and *Aquifex aeolicus* FtsH protein sequences and mapped the available crystal structures onto the sequence alignment (Figure 5A, figure supplement 1). Different transmembrane helix predictors identified both the N-terminal helix and the one linking the periplasmic to the ATPase domain. A region of ~20 aa remained between the second transmembrane helix and the cytoplasmic domain (Figure 5A). Basic Local Alignment Search Tool (BLAST) (Altschul et al., 1997) finds that this 20aa sequence is unique to membrane bound AAA+ proteases. When tested with different structure predictors, this region exhibits extended loop conformation with a weak signal for a short helix at its end (Figure 5, figure supplement 2). Analysis of residue conservation by ConSurf (Ashkenazy et al., 2016) shows 13 highly conserved or conserved residues in this 20 aa region (Figure 5 - supplement 2).

**Figure 5.**
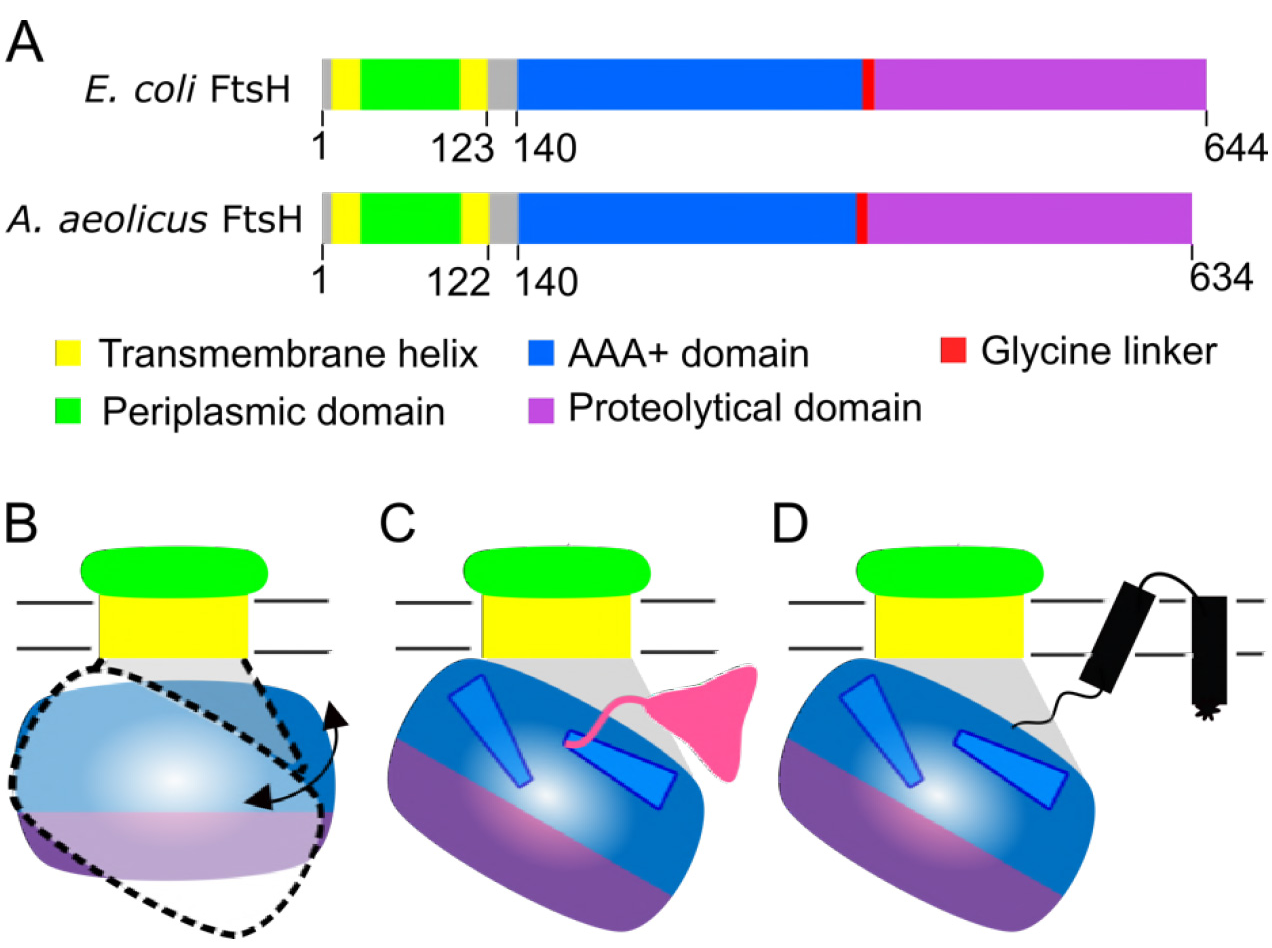
Schematic representation of AaFtsH sequence and proposed model for substrate entry. *A* – Bioinformatics tools and available structures of FtsH domains, from *E. coli* and *A. Aeolicus*, show the presence of a loop-like peptide structure with ~20 aa between membrane and AAA domains. The N-terminal periplasmic domain (green) is between two transmembrane helices (yellow). The second transmembrane helix is followed by a loop region (grey; see text), which is the link to the AAA+ domain (blue). Connected by the glycine linker (red), the C-terminal protease domain is shown in purple. *B* - A new model for substrate entry is proposed. The ~20 aa flexible linker could allow the cytoplasmic domain of FtsH to tilt in relation to the membrane, creating a 30 Å wide gap that provides access of cytoplasmic (pink, *C*) and membrane protein substrates (black, *D*) to the central pore in a partially folded state.

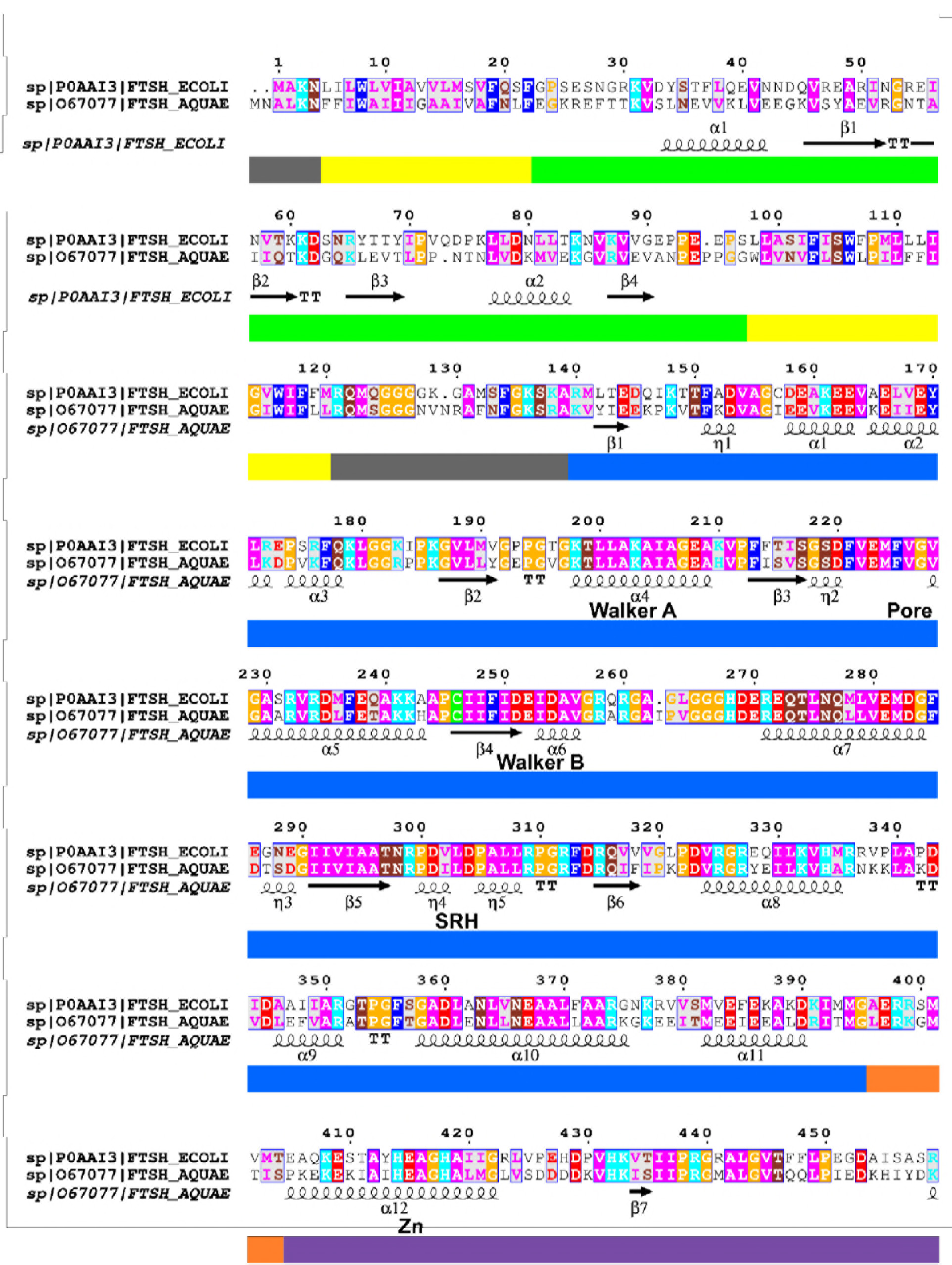

**Figure 5 – figure supplement 1.**
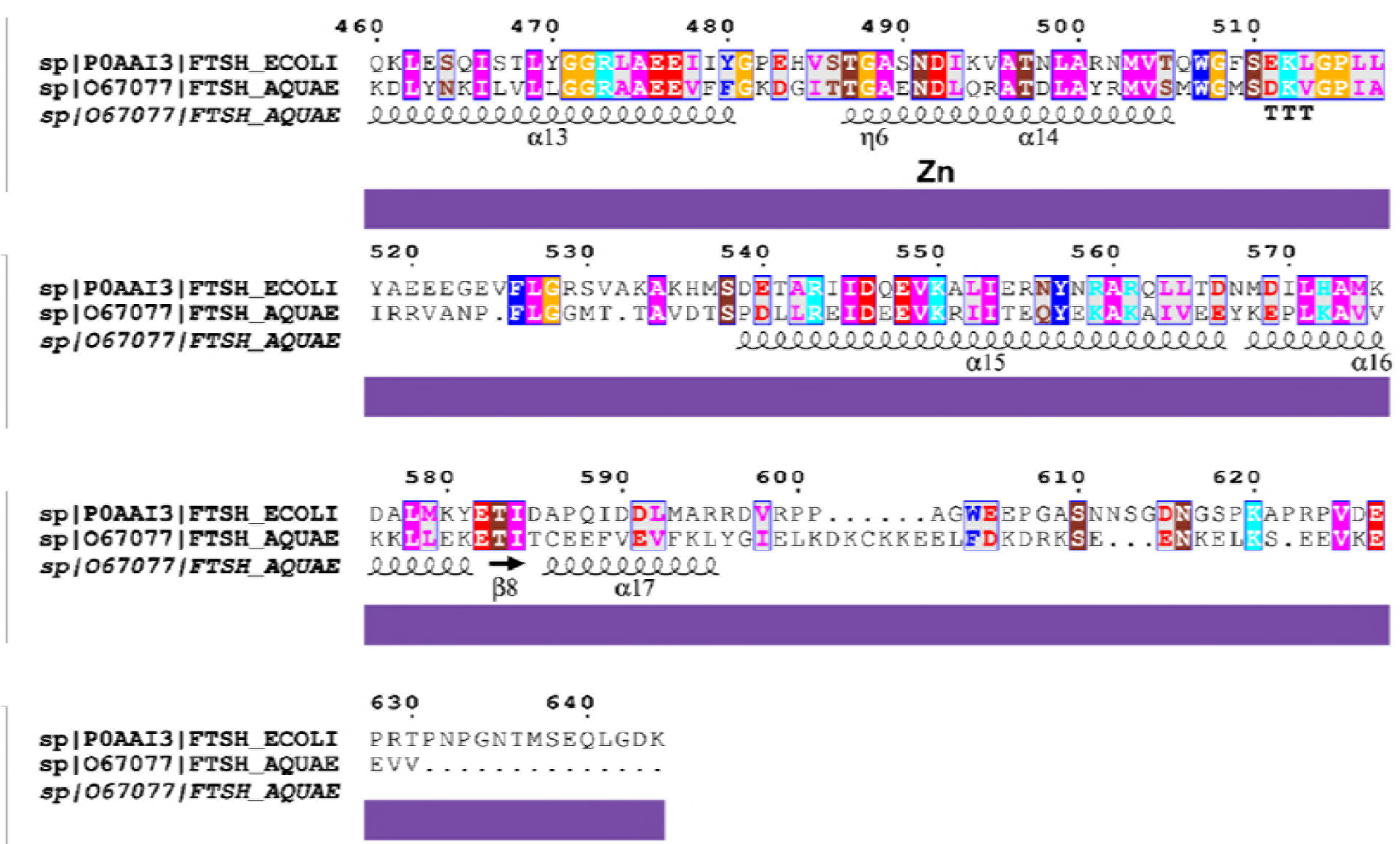
Sequence alignment of the *E. coli* and *Aquifex aeolicus* FtsH full-length sequence. Sequences in crystal structures of Aquifex aeolicus cytoplasm domain (PDB 4WW0) are underlined with coloured bars (AAA domain: blue; glycine linker: red; proteolytic domain: purple), and in the periplasmic domain (PDB 4V0B) with a green bar. Secondary structures are indicated by symbols below the sequence. Yellow bars refer to the two transmembrane helices and the grey bars correspond to the loop-like regions. Transmembrane helix modelling programs (RHYTHM, CCTOP, TMHHM, HMMTOP, TOPCONS and TMpred) predict the N-terminal helix (TM1) to contain 18 residues (FFIWAIIIGAAIVAFNLF) and the second helix (TM2) to contain 23 residues (WLVNVFLSWLPILFFIGIWIFLL). The alignment was performed using ESPript 3 (Grosjean et al., 2010).

**Figure 5 – figure supplement 2.**
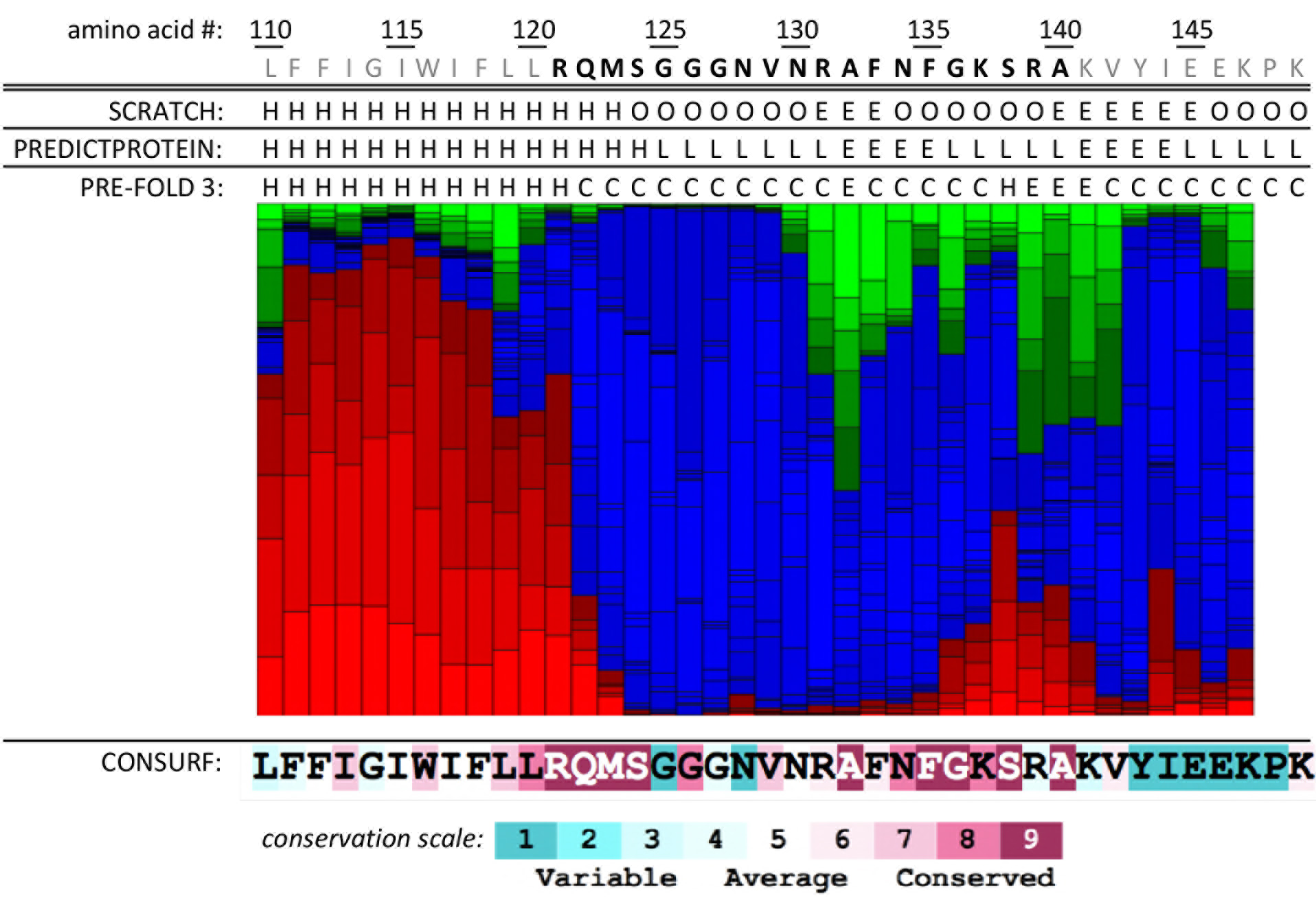
Structure of the ~20 aa region (amino acids in black) between the TM2 and the AAA-domain of *Aquifex aeolicus* FtsH, as predicted by the structure predictors SCRATCH, PREDICTPROTEIN and PRE-FOLD 3 (H: helical; E: extended; L: loop; C: coil; O: other). Residue conservation scores are obtained from the ConSurf server (scale at the bottom).

## Discussion

In this work, we have purified full-length *Aquifex aeolicus* FtsH and characterized both its structure with negative stain EM and its ATPase and proteolytic activities. The purification of full-length FtsH hexamers proved to be challenging with size exclusion chromatograms reproducibly showing two species of AaFtsH in closely overlapping fractions. Negative stain EM and image analysis confirmed that the low molecular weight fraction corresponds to AaFtsH hexamers, and the high molecular weight fraction to AaFtsH dodecamers (Figures 2 and 3). Importantly, both fractions show highly similar specific ATPase activities: 340 nmol/min/mg for AaFtsH dodecamers and 338 nmol/min/mg for AaFtsH hexamers. This compares well to values reported for the *E. coli* FtsH, 230 nmol/min/mg (Tomoyasu et al., 1995) and 193 nmol/min/mg (Bruckner et al., 2003). Also, the Michaelis Menten constants are similar for the dodecameric and hexameric forms, with K_M_ (518 and 600 μM, respectively) and K_cat_ (61 and 63 min^−1^, respectively). The K_M_ values obtained for ATPase activity of hexamers and dodecamers are between 7x (Tomoyasu et al., 1995) and 20x (Bruckner et al., 2003) higher than reported for *E. coli* FtsH solubilised in a different detergent and under different conditions. Both fractions of LMNG-solubilized AaFtsH can degrade a casein substrate at comparable proteolytic rates (Figure 4C, D). Our results also document that AaFtsH hexamer and dodecamer both show high proteolytic activity, since the same amount of casein can be degraded by 10x less LMNG-AaFtsH when compared with AaFtsH solubilized in n-Dodecyl-β-D-Maltoside (DDM) (Vostrukhina et al., 2015). In summary, we demonstrate that the AaFtsH kept in solution by LMNG exists as equally active dodecamers and hexamers.

Molecular weight estimates from SEC suggest that for the hexamer, the LMNG micelle and the bound lipid molecules account together for ~300 kDa (Figure 1). Combined SEC and MALDI-TOF MS measurements showed that the dimeric ABC transporter BmrA, comprised of 12 transmembrane helices modelled as a cylinder of 40 Å diameter, binds an LMNG micelle of 157 kDa (Chaptal et al., 2017). In AaFtsH hexamers, the LMNG micelle containing the 12-helix ring has a width of 100 Å (Figure 3). According to previous reports the LMNG belt is ~20 Å thick (Chaptal et al., 2017; Huynh et al., 2014; Vahedi-Faridi et al., 2013). Therefore, the cylindric transmembrane domain of the hexamer has a diameter of ~60 Å, accommodating 1.5 × 157 or 240 kDa of LMNG. The ring of twelve tilted helices emanating from the periplasmic domain (PDB 4V0B) also exhibits a diameter of ~60 Å, consistent with the above estimate. Transmembrane helix 1 (TM1) corresponds in length to TM4 of aquaporin 1 (AQP1; PDB 1FQY) (Murata et al., 2000) and is likely tilted by 29°. TM2 corresponds to TM6 of AQP1 and is thus probably tilted by 36°. Modelling helices as 10 Å wide tilted cylinders, the cross section of the helical bundle is 6 × π × 5^2^ (1/cos (29°) + 1/ cos(36°)) Å^2^, and the bilayer cross section inside the ring then amounts to 1700 Å^2^. Taking 50 Å^2^ per lipid molecule into account, the ring houses about 34 lipid pairs, which make ~50 kDa. From the dimension of the membrane domain with its LMNG belt measured by EM, the LMNG micelle and the lipid molecules are estimated to represent ~290kDa, close to the molecular weight estimate from SEC. Such a molecular weight estimate is less accurate for the AaFtsH dodecamer, because it elutes outside the range of calibration proteins used. In addition, the hexamer has possibly different migration properties compared to the compact elongated dodecamer composed of two intertwined hexamers (Figure 3).

Further we report here the first full length bacterial FtsH 3D map at a resolution of 20 Å, which is related to the negative staining used for visualizing the protein complexes, as well as their intrinsic flexibility. As documented in Figure 3G this map accommodates X-ray structures of cytosolic and periplasmic fragments. The density of the X-ray structure of *Aquifex aeolicus* cytosolic domain (PDB 4WW0) rendered at 20 Å resolution fits well with the cytosolic domain of the 3D map presented here. There is also a similar match of the periplasmic domain with the periplasmic crystal structure of *E. coli* FtsH (PDB 4V0B). The reported intermembrane domain of *m*-AAA is larger (Lee et al., 2011), as it comprises ~25% more amino acids than the AaFtsH periplasmic domain. Nevertheless, except for the periplasmic domain, the yeast full-length *m-* AAA cryoEM map rendered at 20 Å also firs well with the 3D map of the AaFtsH hexamer.

The unexpected AaFtsH dodecamers consistently observed (Figures 1 and 2) exhibit highly variable conformations (Figure 3H and I), with periplasmic domains strongly tilted with respect to cytosolic domains. These dodecamers composed of intertwined hexamers that share a single LMNG micelle reveal a large flexibility of the region that links the membrane and ATPase domains. While the 12 Å cryo-EM structure of *m*-AAA shows these thin links clearly (Lee et al., 2011), the subsequent model of the full-length *E. coli* FtsH predicts transmembrane helices projecting into the ATPase domain (Scharfenberg et al., 2015), which would not allow the conformational flexibility we observe. However, this model is neither compatible with the thin links shown by Lee et al., 2011, nor with the significant stain-penetrated gap between the cytosolic AaFtsH domain and membrane domain visualized in Figure 3.

Based on these results we propose a new model for substrate entry. Conformational variations demonstrated by the AaFtsH dodecamers (Figure 3) indicate that FtsH is flexible enough for the large cytosolic domain to tilt significantly with respect to the membrane plane, thus accommodating larger substrates (Figure 5B-D). In the fully active dodecamer, the space between the cytosolic domain and the membrane compares to the height of the AaFtsH periplasmic domain. Thus, our results suggest that the gap at the edge of the cytosolic domain of the native, hexameric AaFtsH may open up to ~30 Å. Such a gap is enabled by the ~20aa linker present between the second transmembrane helix and the cytosolic domain sequence (Figure 5A, grey). This linker appears to have remarkable properties: it exhibits a length of ~13 Å in its quiescent state, but can extend like a spring at least up to ~30 Å. Structure predictors produce similar models that include a possible elongation of TM2, comprising at most the highly conserved RQM(S) motif, followed by a loop (GGGNVN), and a conserved region that may adopt loop, extended strand or helical configurations (RAFNFGKSRA) (Figure 5 - supplement 2). This linker’s possible extended loop conformation explains the structural variation of the dodecameric form (Figure 3H-K), and would also accommodate the large movements of the ATPase domain predicted by X-ray crystallography (Bieniossek et al., 2009). Endpoints of 20 aa loops are separated by a minimum distance of ~10 Å in known proteins structures, but can extend or be stretched up to a maximum of 60-80 Å (Ainavarapu et al., 2007; Choi et al., 2013). These large tilting movements of the cytosolic domain of FtsH would allow misfolded proteins to access the pore (Figure 5 B-D), without the need for an unfolded polypeptide to snake through a 13 Å channel before reaching to translocating loops (Lee et al., 2011).

Although we do not anticipate FtsH to form dodecamers in bacteria, we propose that the large tilting movements observed here are likely achieved during proteolytic activity *in vivo*. A higher resolution model of full-length FtsH would give us more details on how FtsH interacts with its substrates. This has however proven to be difficult, precisely due to the high flexibility of the 20 aa linker at the membrane junction.

## Materials and Methods

### AaFtsH expression and purification

Full-length *Aquifex aeolicus* FtsH (AaFtsH) cloned into pET22a vector was kindly granted by Ulrich Baumann and the same expression conditions were employed (Vostrukhina et al. 2015). Cells were harvested at 3500 *g* for 25 min at 4 °C and disrupted in a cell disruptor. The resulting cell debris was purified at 20000 *g* for 15 min and membranes were isolated at 125000 *g* for 3 hours. Membranes were solubilised in 20 mM Tris-HCl pH 8.0; 150 mM NaCl; 1%(w/v) Lauryl Maltose Neopentyl Glycol (LMNG) (Anatrace) for 3 hours at 4 °C and cleared at 125000 *g* for 1 hour at 4 °C. The sample was first purified by affinity chromatography, using a HisTrap-5 mL column (GE Healthcare). FtsH fractions were eluted in 20 mM Tris-HCl pH 8.0; 500 mM NaCl; 0.01%(w/v) LMNG and 200 mM imidazole. AaFtsH fractions were then incubated at 60 °C overnight with 20 mM ATP; 10 mM MgCl_2_; and 25 μM ZnCl_2_. The incubated sample was concentrated to 500 μL and loaded into a SEC Superose 6 Increase 10/300 GL column (GE Healthcare) pre-equilibrated with 10 mM Tris-HCl pH8.0; 150 mM NaCl; 0.01%(w/v) LMNG; 5% Glycerol. AaFtsH monomer concentration was measured with a Nanodrop. A representative SEC run is plotted in Figure 1A. The centre position of the two largest peaks was determined by fitting Gaussian functions to 10 SEC profiles. A calibration curve of the Superose 6 Increase 10/300 GL was performed using the Gel Filtration High Molecular Weight Calibration Kit (GE Healthcare), following the GE Healthcare instructions. AaFtsH fractions were analysed by Native PAGE gels using the MiniPROTEN^®^TGX™ Precast Protein Gels (Biorad).

### Transmission Electron Microscopy analysis

3 μL of AaFtsH dodecamer or hexamer fractions was loaded on a carbon-coated 400 square mesh copper grid (Aurion) previously glow-discharged for 1 min. The liquid drop was absorbed with filter paper after 1 min and quickly washed with a drop of water that was again blotted with filter paper. This procedure was repeated 3 times to rinse all the detergent present in the samples. Finally, a 3 μL drop of 3 % uranyl-acetate was added to the grid, incubated for 1 min and absorbed with a filter paper. Transmission electron microscopy was performed using a Philips CM-200T and a JEOL 3200 FSC both equipped with a TemCam-F416 (TVIPS) and recorded at 50000x magnification using the EM-MENU software with a sampling rate of 2.23 Å/pixel.

### ATPase activity

The ATPase activity was measured using the High Throughput Colorimetric ATPase Assays kit (Innova Biosciences). Free phosphate from the AaFtsH dodecamers and hexamers was eliminated by an incubation with 100 μL of PiBind™ resin (Innova Bioscience) for 30 min. Also mix A [50 mM Tris-HCl pH 8.0; 150 mM NaCl; 10 mM MgCl_2_], was previously incubated with PiBind™ resin (Innova Bioscience). Reactions were started by mixing AaFtsH (0.25 μM final concentration) in mix A with either 50, 100, 250, 500, 1000, or 1800 μM ATP (final concentrations). Reaction rates were measured every 2 min for a total of 10 min, in triplicates. ATP concentrations were chosen such that the maximum concentrations were more than 10x the previously reported K_M_ values (Bruckner et al., 2003; Tomoyasu et al., 1995). 25μL of P_i_ColorLock™ mix was added to stop the reaction, and after 5 min, 10 μL of stabiliser reagent was added.

OD_650_ was measured after 30 min using a CLARIOstar (BMG-Labtech) microplate reader, every replica was measured 3 times and an average of these readings was calculated for each replica (Figure 4 - figure supplement 1). Released Pi concentrations were calculated from a calibration curve of standard Pi concentrations measured. K_M_ and K_cat_ were calculated for both hexamers and dodecamers. Michaelis-Menten constants were obtained, assuming the steady-state of the reaction, with GraphPad Prism software with a 95% Confidence Interval (CI).

### Protease activity

AaFtsH protease activity was assessed using the Resorufin labelled casein kit (Roche). 0.25, 0.5, 1.0 or 2.0 μM (final concentrations) of AaFtsH dodecamers or hexamers were incubated with 50 μM of resorufin labelled casein in [50 mM Tris-HCl pH 8.0; 80 mM NaCl; 12.5 μM ZnCl_2_; 5 mM MgCl_2_; 1 mM dithiothreitol; 0.01% LMNG and 10 mM ATP] at 60 °C. Triplicates of these measurements were taken. Reaction rates were measured every 10 min for 30 min (Figure 4 - figure supplement 2). 160 μL of 5% trichloroacetic acid was added and incubated at 37 °C for 30 min. Proteins were precipitated at 16100 *g* for 30 min and 120 μL from the supernatant was mixed with 80 μL of 500 mM Tris-HCl pH 8.8 and added to a 96 well transparent plate. The OD_574nm_ was immediately measured using an Infinite^®^ 200PRO (TECAN) plate reader.

### Sequence alignment and structure prediction

The AaFtsH and *E. coli* FtsH sequences were aligned with ESPript 3 (Grosjean et al., 2010). Part of the AaFtsH sequence (1-144 aa) sequence was submitted to different transmembrane helices structure predictors: RHYTHM (Rose et al., 2009), CCTOP (Dobson et al., 2015), TMHHM (Krogh et al., 2001), HMMTOP (Tusnády and Simon, 2001), TOPCONS (Tsirigos et al., 2015) and TMpred (Expasy) (Hofmann and Stoffel, 1993). Prediction of the transmembrane helices was performed to identify the beginning of the linker between membrane and AAA domains. To explore whether this 20 aa linker has a known structure, part of the sequence (110-148) was submitted to the structure predictors SCRATCH (Cheng et al., 2005), PRE-FOLD 3 (Lamiable et al., 2016) and PREDICTPROTEIN (Rost et al., 2004). Sequence similarity search was performed with BLAST (blast.ncbi.nlm.nih.gov) (Altschul et al., 1997). Residue conservation was assessed with the ConSurf server (Ashkenazy et al., 2016).

### Imaging Processing of negatively stained single particles

Single particle imaging processing was performed using Scipion1.1 (Rosa-Treivn de la et al., 2016) and Eman2.12 software packages (Ludtke,2016). A set of 15000 particles was manually picked using a semi-automated mode, the particles were 2D averaged into 100 classes. To measure the dimensions of the negatively stain particles two approaches were taken. First, the height of dodecamers and hexamers was measured directly from 280 particles in the micrographs. Measurements referring to the different domains were done on 10 class averages chosen. The resolution of the maps was calculated from the Fourier Shelf Correlation (FSC) values as implemented in Eman2.12. Using Chimera4.0.4 (Pettersen et al., 2004) different crystal structures of the cytoplasmic domain *Aquifex aeolicus* (PDB 4WW0), the CryoEM of the full-length *m*-AAA (EMDB 1712) and the crystal structure of the *E. coli* periplasmic domain (PDB 4V0B) were automatically fitted into our 3D map.

## Acknowledgements

The authors thank Dr. Simon Lindhoud for useful initial discussions; Prof. Dr. Ulrich Baumann for the plasmids; and Dr. Carlos Oscar Sorzano, Prof. Jose Maria Carazo and Prof. Steven Ludtke for useful discussions of single particle imaging processing. The work was supported by the Netherlands Organization for Scientific Research (VIDI NWO Grant 723-016-007).

